# AlphaFold2 has more to learn about protein energy landscapes

**DOI:** 10.1101/2023.12.12.571380

**Authors:** Devlina Chakravarty, Joseph W. Schafer, Ethan A. Chen, Joseph R. Thole, Lauren L. Porter

## Abstract

Recent work suggests that AlphaFold2 (AF2)–a deep learning-based model that can accurately infer protein structure from sequence–may discern important features of folded protein energy landscapes, defined by the diversity and frequency of different conformations in the folded state. Here, we test the limits of its predictive power on fold-switching proteins, which assume two structures with regions of distinct secondary and/or tertiary structure. Using several implementations of AF2, including two published enhanced sampling approaches, we generated >280,000 models of 93 fold-switching proteins whose experimentally determined conformations were likely in AF2’s training set. Combining all models, AF2 predicted fold switching with a modest success rate of ∼25%, indicating that it does not readily sample both experimentally characterized conformations of most fold switchers. Further, AF2’s confidence metrics selected against models consistent with experimentally determined fold-switching conformations in favor of inconsistent models. Accordingly, these confidence metrics–though suggested to evaluate protein energetics reliably–did not discriminate between low and high energy states of fold-switching proteins. We then evaluated AF2’s performance on seven fold-switching proteins outside of its training set, generating >159,000 models in total. Fold switching was accurately predicted in one of seven targets with moderate confidence. Further, AF2 demonstrated no ability to predict alternative conformations of two newly discovered targets without homologs in the set of 93 fold switchers. These results indicate that AF2 has more to learn about the underlying energetics of protein ensembles and highlight the need for further developments of methods that readily predict multiple protein conformations.

## Introduction

Deep learning-based algorithms have made it possible to predict protein structure from amino acid sequence, sometimes with impressively high accuracy. The most successful of these algorithms, AlphaFold2 (AF2) (1), has inspired numerous new approaches to predict and design other important structural features of proteins. These features include protein-protein interaction sites (2), conditionally folding regions of intrinsically disordered proteins (3), and structures of novel protein folds from metagenomic sequences (4).

The many successes of AF2 suggest that it may also be able to predict subtle-yet-important protein properties previously revealed only through sophisticated techniques. These properties include conformational ensembles and functionally important alternative conformations (5). Consistent and accurate predictions of these properties would suggest that AF2 may do more than simply associate protein sequence with structure through sophisticated pattern recognition (6). Rather, it may leverage learned folding physics to accurately approximate folded protein energy landscapes (7). These landscapes are defined by the diversity and frequency of protein conformations in the folded state. Supporting this possibility, AlphaFold2 has successfully predicted alternatively folded states in over a dozen cases (5, 8).

Yet despite AF2’s impressive accuracy and broad success, several uncertainties remain about how much it has learned about protein energy landscapes, particularly its ability to predict alternative protein conformations. These uncertainties relate to the two major tasks on which protein structure prediction relies: adequate sampling and accurate scoring. First, sampling refers to AF2’s ability to generate distinct experimentally consistent conformations of the same protein. As a deep learning algorithm, AF2 relies on a large training set of hundreds of thousands of solved and predicted structures, their amino acid sequences, and multiple sequence alignments (MSAs) containing the evolutionary information used to infer structure (1). Compared to this large training set, the number of experimentally determined alternative protein conformations is small (9). Furthermore, AF2’s ability to sample alternative conformations has been tested on only a handful of examples (5, 8). Thus, it is unknown how well AF2 would accurately sample experimentally consistent alternative protein conformations more broadly (9). Second, scoring refers to AF2’s ability to distinguish between good and poor predictions. Currently, AF2 assigns good and poor scores to its predictions of single protein conformations very reliably (7). To our knowledge, however, no studies have systematically assessed how accurately it scores alternative protein conformations.

Here, we systematically assess AF2’s ability to sample and score both experimentally determined conformations of 93 fold-switching proteins (10). This newly emerging class of proteins has been evolutionarily selected to assume two distinctly folded states (11), presumably for functionally important reasons (12). Previously, we showed that AF2.2.0 is systematically biased to predict one conformation of fold switchers while missing the other (13). Since then, AF2.3.1 has been released: this version now makes accurate predictions of oligomeric assemblies and protein-protein interactions (14). Because at least one conformation of most fold switchers forms an oligomer or interacts with another protein (10), we aimed to assess AF2.3.1’s ability to predict fold switching when information about oligomeric state or binding partner is provided. Both conformations of all 93 fold switchers were deposited in the Protein Data Bank (15) (PDB) before AF2.3.1 was trained, suggesting that they are likely in its training set. Furthermore, two methods for predicting alternative protein conformations or protein ensembles with AF2 have recently been proposed (16, 17). Thus, we tested the performance of these methods on the same set of 93 fold switchers, generating >280,000 predictions in all. Upon assessing these predictions, we found that AF2 predicts fold-switching proteins likely in its training set with modest success (23/93). Further, its confidence metrics select against alternatively folded protein conformations and cannot discriminate between low and high energy conformations of fold-switching proteins. Because AF2’s predictions are most useful for proteins without experimentally determined structures, we also tested it on a set of seven fold-switching proteins whose structures were not deposited in the PDB at the time AF2.3.1 was trained, generating >160,000 additional predictions. It failed to predict the alternative folds of 6/7 fold switchers. These results call into question how much AF2 has learned about protein folding energetics and indicate the need for further developments of methods that accurately predict alternative protein conformations.

## Results

### AF2 samples both conformations of ∼25% of fold switchers

AF2’s ability to sample two folds assumed by single sequences was tested on 93 pairs of experimentally determined fold switchers. The structures of all 93 pairs (**Table S1**) are from many diverse fold families and source organisms (13). All structures were deposited in the PDB before 2022 and are therefore likely in AF2.3.1’s training set. All protein pairs have identical or nearly identical sequences and regions of distinct secondary and tertiary structure (**Methods**). AF2 predictions are defined as successful when they accurately capture both experimentally determined conformations, called Fold1 and Fold2. Prediction accuracy is assessed by calculating the TM-score (18) between each AF2 prediction and both experimentally determined structures. TM-scores quantify the similarity of topology and connections between secondary structure elements (19), a reliable metric since fold-switching proteins are identified by secondary structure differences (10). Because whole-protein TM-scores often overestimate the prediction accuracies of fold-switching regions, we assessed predictions using TM-scores of fold-switching regions only (**Figure S1**). Higher TM-scores indicate predictions closer to experimentally determined conformations. We ordered each pair of fold switchers so that Fold1 corresponds to the target conformation most frequently predicted by AF2, and Fold2 corresponds to the less frequently predicted target conformation (**Methods**).

First, four different AF2.3.1 modes were tested on each fold-switching sequence: with templates, without templates, multimer model on single chains, and multimer model on protein complexes (**Table S2**). AF2.3.1’s performance increased slightly above AF2.0’s (**Figure 1A**), capturing 11/93 fold switchers (combining results both with and without templates) rather than 8/93 (13). Furthermore, AF2_multimer successfully predicted both conformations of 12/93 fold switchers. Although fold switching is often triggered by protein-protein interactions (10), supplying information about binding partners and oligomeric states to the Multimer model yielded only four fold-switch predictions, all of which were predicted using single chains by other AF2.3.1 methods (**Table S2**). To augment this TM-score-based assessment, we also performed root-mean-square-deviation (RMSD) calculations of fold-switching regions and found similar results (**Figure S2**). Together, these assessments demonstrate that running AF2.3.1 with default inputs and parameters rarely produces successful fold switch predictions: <13% in total.

**Figure 1.**
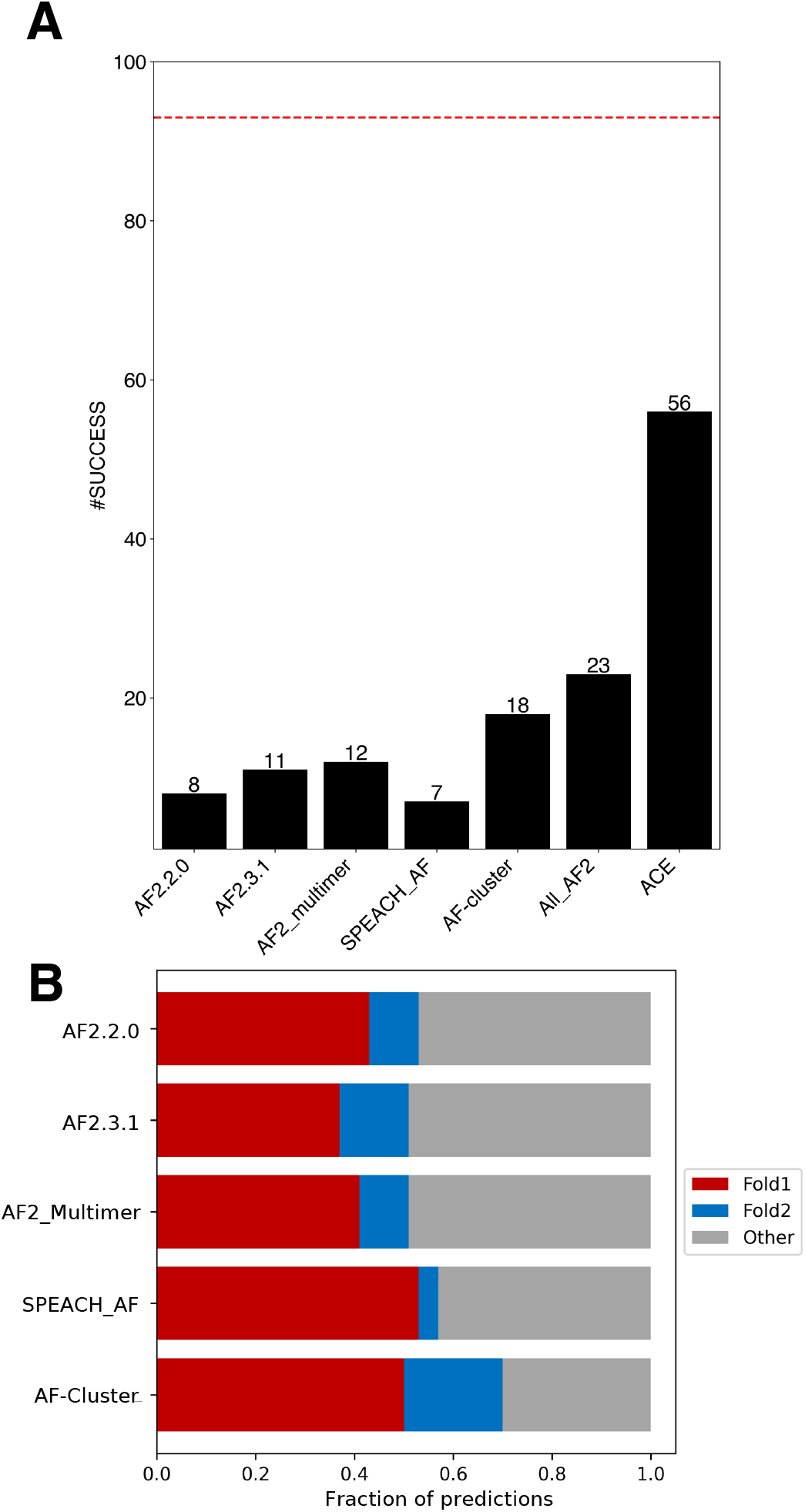
AF2 predicts fold switching with modest success. (a) Numbers of successful fold-switch predictions for each AF2 method compared with coevolutionary information found for both folds (ACE) and the total number of possible successes (dotted red line). All_AF2 combines all unique successful predictions from all AF2-based methods: >282,000 predictions. (b). Fraction of predicted structures that match experimentally determined conformations for all methods. Fold1 is the conformation most frequently sampled by AF2.3.1, Fold2 is the less frequently sampled (or unsampled) conformation. Conformations designated as Other are inconsistent with both experimentally determined structures.

We then tested whether AF2-based enhanced sampling approaches can predict more fold switchers than AF2 runs with standard inputs. Recently, two such approaches have been proposed to predict alternative conformations of proteins including fold switchers. The first, SPEACH_AF (16), masks coevolutionary information in AF2’s input MSA by mutating selected columns to alanine *in silico*. Masking this information is expected to allow AF2 to identify coevolutionary signals in the MSA corresponding to alternative protein conformations, allowing it to sample a more diverse conformational ensemble. SPEACH_AF was tested on 16 different proteins and generated alternative conformations for almost all of them. Though none of these proteins were fold switchers, SPEACH_AF’s potential to predict fold switching was proposed (16). The second approach, AF-cluster (17), clusters sequences from a deep MSA by similarity and runs AF2 on different clusters or combinations thereof. This approach is based on the hypothesis that different MSA subsets may contain coevolutionary information distinct from deep MSAs, allowing AF2 to predict alternative protein conformations, though recent work suggests that AF-cluster infers alternative conformations from sequence similarity rather than coevolution (20). Regardless, AF-cluster was tested on six families of fold-switching proteins and successfully predicted both conformations in three families (17).

To gauge how frequently SPEACH_AF and AF-cluster predict fold switching, we tested both approaches extensively on the set of 93 fold switchers tested previously, generating >77,000 structures with SPEACH_AF and >200,000 structures with AF-cluster (**Table S2**). Both methods missed fold switching in most cases (**Figure 1A**): 94% for SPEACH_AF (6/93 successes) and 81% for AF-cluster (18/93 successes). Interestingly, the alternative folds of 6/17 AF-cluster predictions were also correctly predicted from single sequences. Because AF2 requires an input MSA to make coevolutionary inferences, these single-sequence predictions indicate that AF-cluster’s predictive success did not result from coevolutionary inference in these cases and may have arisen from overtraining instead ((9, 20, 21) **Table S2, Figure S3**).

As mentioned previously, both SPEACH_AF and AF-cluster postulate that AF2 can predict alternative protein conformations when sufficient coevolutionary information is provided. A recent computational approach called Alternative Contact Enhancement (ACE) identified coevolutionary information unique to both folds of 56 fold-switching proteins, confirming that MSAs often contain structural information unique to both conformations (11). Nevertheless, after combining all correctly predicted fold switch pairs from 282,000 predicted structures (**Figure 1B**), AlphaFold2 misses this information in 43/56 cases. Thus, current enhanced sampling approaches typically do not enable AF2 to consistently detect the dual-fold coevolutionary information present in many MSAs of fold-switching proteins.

### AF2 confidence metrics select against alternative conformations of fold switchers

Though AlphaFold2 often produces structural models with remarkably high accuracy, some of its predictions can be inaccurate (1). We quantified the frequency of inaccurate predictions relative to correct predictions of Fold1 and Fold2 generated by all methods (**Figure 1B**). In all cases, 30-49% of predictions did not correspond well to either experimentally determined structure.

To see if AF2 could distinguish between good and inaccurate predictions, the relationship between prediction quality and AF2’s confidence metrics was assessed. AF2 estimates prediction quality with two confidence metrics: the per Residue Local Difference Distance Test (pLDDT) and predicted template modeling (pTM) scores. We sought to determine whether either or both metrics discriminate between the good and poor fold-switch predictions generated by AlphaFold2 and AF-cluster. AF-cluster was selected because it predicted substantially more fold switchers than SPEACH_AF (18 rather than 7), generated fewer inaccurate predictions overall (∼30% rather than 43%), and comprised a larger set of predictions.

Neither of AF2’s confidence metrics successfully discriminated between good and inaccurate fold-switch predictions (**Figures 2A, S4-S6**). Rather, both pLDDT and predicted template modeling (pTM) scores assigned lower confidences to diverse correctly predicted conformers and higher confidences to incorrect predictions. Whereas 30% of all AF-cluster structures did not match experimentally determined structures of Fold1 or Fold2–the fewest incorrect predictions of all methods (**Figure 1b**)–nearly 70% of the highest ranked structures were inconsistent with experiment (**Figures 2A, S4, Table S3**). A similar trend was observed for AF2.3.1 runs with standard settings (**Figures S5, S6**).

**Figure 2.**
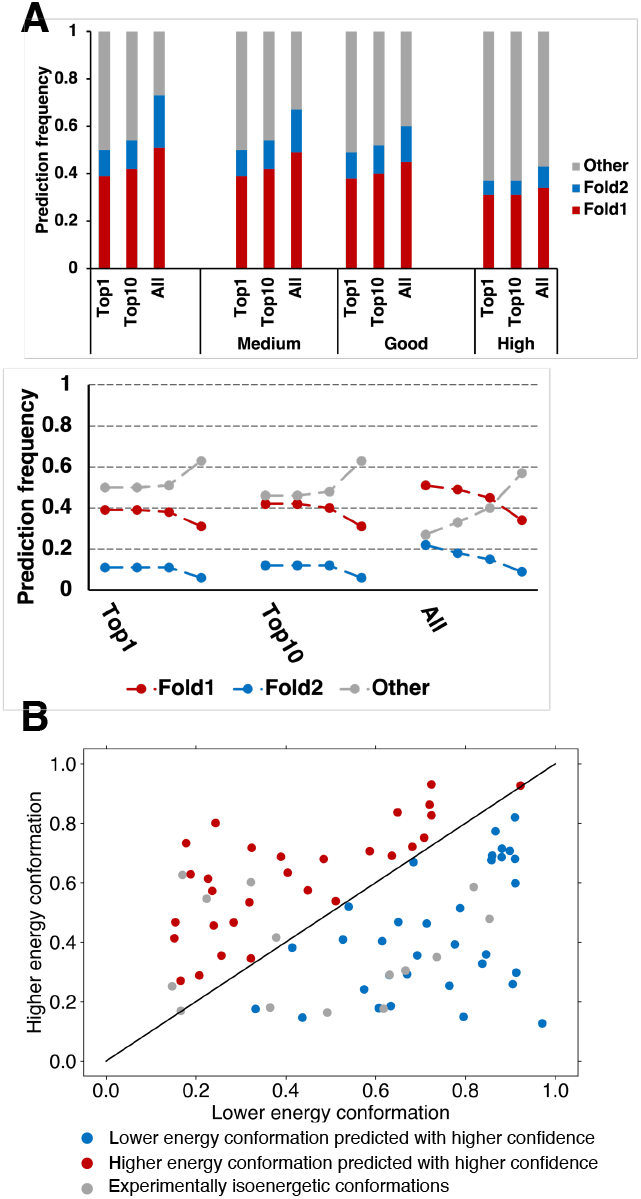
AF2 confidence metrics select against alternative conformations and do not predict the most energetically favorable fold-switch conformations. (A) Bar-plot representation of prediction success in Top1, Top10 and All fold-switch predictions select more incorrect conformations as prediction confidence increases. These trends are apparent in trendline plots showing the change in fraction of predictions as a function of prediction confidence. The leftmost 3 trendlines are from All predictions, the middle/rightmost are from Top10/Top1 most confident for each of 93 fold switchers. For each column of trendlines, the leftmost dot represents all conformations (not weighted by confidence), the next is predictions with medium confidence, then good confidence, and finally high confidence. Confidences are determined by ≥70% (medium), 80% (good), 90% (high) of residues with pLDDT scores ≥70. (B). AF2’s structure module predicts the lower energy conformations of fold switchers with better accuracy and higher confidence than higher energy conformations 50% of the time, equal to random chance. Blue dots represent correctly predicted ground state conformers with higher confidence; red dots represent correctly predicted excited state conformers with higher confidence than low energy, and gray dots have been observed to sample both folds at roughly equal proportions at equilibrium. Axes represent TM-scores of both conformation relative to experiment.

These results strongly indicate that AF2’s confidence metrics select against experimentally consistent predictions of fold switchers, especially Fold2, in favor of experimentally inconsistent predictions. For instance, while AF-cluster correctly predicted 19/93 Fold2 conformations overall, only 7/93 were identified amongst high quality predictions (p < 8.1*10^-4^, one-sided binomial test). Further, significantly fewer correctly predicted conformations (either Fold1 or Fold2) were identified amongst high-quality models (37) than amongst all (53, p < 6.6*10^-4^, one-sided binomial test). Some of these experimentally unobserved conformations have been proposed to correspond to folding intermediates (5). To the best of our knowledge, there is no experimental evidence supporting this claim for fold-switching proteins. In fact, a recently characterized folding intermediate of the transcriptional regulator RfaH suggests the opposite (22). AF2-multimer predicted a hybrid α-helical/β-sheet fold with high confidence for its fold-switching C-terminal domain (**Figure S7**). This prediction is not consistent with experiment: most notably, the N-terminal portion of the AF2 prediction folds into a β-hairpin, while the experimentally observed intermediate has helical propensities in that region (22). Thus, high confidence AF2 predictions that differ from experimentally determined structures do not necessarily correspond to folding intermediates, consistent with previous observations (23).

AF2’s inability to discriminate between good and poor predictions of fold switchers suggests that its confidence metrics may have broader limitations. To further assess this possibility, we used AF2’s structure module to energetically rank fold-switching protein pairs (**Methods**). This approach–based on hypothesis that the AF2’s structure module has learned protein folding physics–correctly selected experimentally consistent structures among diverse models of 283 proteins (7). Here, it correctly selected the ground state conformations of fold-switching proteins 50% of the time (**Figure 2B, Methods**). In other words, the selective power of AF2’s structure module was tantamount to random guessing for fold-switching proteins. It may seem reasonable to hypothesize that this selective failure arises in cases where the ground states of fold switchers are oligomeric and the excited states are monomeric. This may not be the case, however, because AF2 predicts the folds of ground state oligomeric structures, such as KaiB, with the monomer model (17). Furthermore, including oligomeric states and binding partners in the multimer model did not produce any new fold-switch predictions (**Table S2**); instead, all alternative conformations were predicted from monomeric sequences without the need for additional information about oligomeric state or binding partner. Thus, AF2 can correctly predict the conformations of single chains of homo- or hetero-oligomers without additional information about oligomeric state or protein binding partner.

### AF2 rarely predicts newly identified fold switchers

AF2’s modest success in sampling the conformations of fold switchers likely within its training set raises the question of how well it can predict fold switching of sequences without. After all, AF2 is most valuable when used to infer structural properties of uncharacterized proteins, such as conditionally folding regions of IDPs (3) and novel folds (4). Thus, we identified seven fold switchers with sequences outside of AF2’s training set and divided them into two categories: distant homologs of a known fold switcher and newly discovered fold switchers. The alternative conformations of all seven fold switchers were either (1) determined after AF2.3.1’s last training or (2) inferred by other experimental methods without depositing the alternative structure in the PDB.

First, we assessed AF2’s ability to predict fold switching of five distant homologs of the known fold-switching protein *Escherichia coli* RfaH (24), a bacterial transcription factor whose C-terminal domain reversibly switches from an all α-helical ground state to an all β-sheet excited state upon binding RNA polymerase and a specific DNA sequence called *ops* (25). Both conformations of *E. coli* RfaH have been determined experimentally (26, 27). Previous work provided circular dichroism (CD) and nuclear magnetic resonance (NMR) evidence for switching in all five of these sequence-diverse RfaH homologs (24), all <35% identical to one another and to *E. coli* RfaH. As a control, AF2’s ability to predict single folding was assessed in five additional experimentally characterized single folding RfaH homologs whose CTDs were found to assume the β-sheet fold only (**Table S4**).

Although both AF2 and AF-cluster correctly predict that *E. coli* RfaH–likely in AF2’s training set–switches folds, neither reliably predicted fold switching in the experimentally confirmed variants not deposited in the PDB. Specifically, AF2.3.1 predicted a helical CTD in 1/5 cases with moderate confidence (**Figure S8**). In the other four cases, it predicted the β-sheet conformation only, as it did correctly for all single-folding controls. To extensively search for fold switching with AF-cluster, we generated 50 models per input MSA with 10 seeds for a total of 140,050 predictions of 10 proteins (**Table S5, Methods**). AF-cluster predicted both folds for 4/5 conformations and only well-folded β-sheet conformers in the remaining case (**Figure S9**). All helical conformations were predicted with low confidence (average pLDDT ≤ 50), indicating that AF2.3.1 can generate more trustworthy helical CTD predictions thant AF-cluster. This finding is consistent with the original AF2 paper’s observation that MSAs with ≥32 sequences are needed for reliable predictions (1); AF-cluster-generated MSAs often have ≤10 sequences. Importantly, AF-cluster predicted low-confidence helical conformations in two RfaH homologs with CTDs experimentally confirmed to assume β-sheet folds rather than α-helical (**Figure S9**). NMR evidence from a previous study strongly suggests that the *Candidatus Kryptonium thompsoni* variant assumes the β-sheet conformation only (24). Furthermore, the CD spectrum of the *T. diversioriginum* variant also suggests that it assumes a ground state β-sheet structure consistent with previously characterized RfaH variants whose CTDs do not assume helical conformations (**Figure S10**). Together, these results demonstrate that neither AF2 nor AF-cluster reliably predict fold switching of distant RfaH homologs.

Structures of the two remaining prediction targets were deposited into the PDB in 2023, after AF2.3.1 was trained. Fold switching of Sa1–a 95 amino acid protein that reversibly interconverts between a 3-α-helix bundle and an α*/*β plait fold in response to temperature–was demonstrated by NMR spectroscopy (28). We also included the structure of BCCIPα, a human protein whose sequence is 80% identical to its homolog BCCIPβ. Although BCCIPα has not been shown to switch folds, it assumes a completely different structure than BCCIPβ and has a different binding partner (29). Previous work has shown that when run with default parameters, AlphaFold2 fails to predict the unique structure of BCCIPα, whose most similar PDB analog differs by 9.9Å (29). Thus, we included BCCIPα because (1) we wanted to see if AF-cluster could produce its unique structure and (2) although BCCIPα might not switch folds, it tests AF2’s limits in predicting novel protein folds.

AF-cluster missed fold switching completely for both Sa1 and BCCIPα (**Figure 3**). Specifically, 98.8% (2520/2550) of the Sa1 predictions assumed the α*/*β plait fold, and 54% (2022/3750) of the BCCIPα predictions assumed the structure of its PDB homolog BCCIPβ. By contrast, AF-cluster failed to predict both the 3-α-helix bundle conformation of Sa1 and the experimentally determined conformation of BCCIPα. BCCIPα’s structure was solved in complex with another protein (29). Running the Multimer model with BCCIPα’s binding partner still yielded the BCCIPβ structure (**Figure S11**). Because its apo structure has not yet been determined, it is possible that apo BCCIPα assumes the same structure as BCCIPβ, in which case AF2 and AF-cluster fail to predict its alternative conformation. It is also possible that apo BCCIPα assumes the same structure in its apo and bound forms, in which case AF2 and AF-cluster fail to predict its structure altogether. These results cast doubt on the AF2’s reliability and consistency in predicting the alternative conformations of fold switchers outside of its training set.

**Figure 3.**
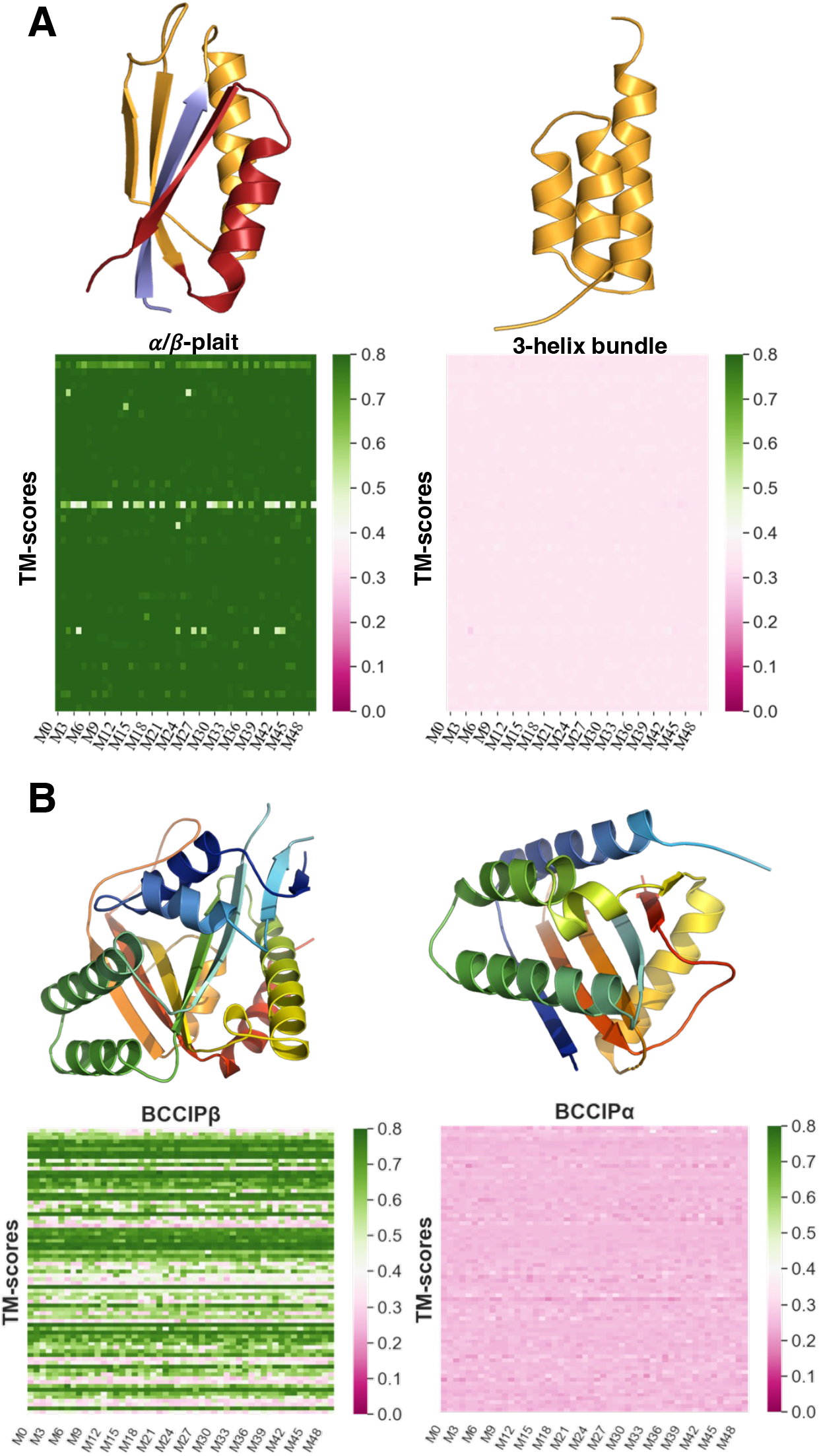
AF2 fails to predict fold switching in two newly discovered cases outside of its training set. (A). Sa1 is a designed protein that switches reversibly between α/β-plait ( PDBID:8e6y, Fold1) and 3α helix (PDBID: 2fs1, Fold2) folds triggered by temperature changes. Cartoon representations of Fold1 are colored blue for N-terminal residues (1 to 10), orange for the fold-switching residues (11 to 66 aligning with the amino acid sequence in Fold2, also in orange) and C-terminal residues (67 to 95) are red. Heatmaps of 50 predictions (M0 to M49) for each of 51 sequence clusters showing the similarity (TM-scores) to Fold1(left panel) and Fold2 (right) are presented below the cartoon representations of the two states. AF-cluster consistently predicts Fold1 but misses Fold2. (B). BCCIPβ and BCCIPα are human protein isoforms with 80% sequence identity that adopt distinct folds. (13Å RMSD). AF-cluster was run on BCCIPα’s sequence. In the right panel, a cartoon representation of BCCIPα (colored blue to red from N-terminus to C-terminus) is shown with the heatmap of TM-scores of 50 predictions (M0 to M49) for each of 75 sequnce clusters compared to the fold adopted by the α isoform (PDBID:8exf, chain B). In the left panel, the BCCIPβ experimental structure (PDBID:7kys) is shown with the heatmap of TM-scores compared to the fold adopted by the β isoform. AF-cluster frequently predicts the structure of the β-isoform but misses the experimentally consistent α-isoform structure.

## Discussion

Although AF2 has revolutionized protein structure prediction and protein design, its current ability to predict alternative protein conformations is limited. We tested multiple versions of AF2 and two published enhanced sampling methods on 93 fold-switching proteins, which assume two distinct biologically important conformations (10, 12). Although both conformations of all 93 fold switchers were likely in the latest version of AF2’s training set, combining all models from all methods and sampling techniques–>280,000 predictions in all–captured fold switching with a modest success rate of 24% (23/93). For context, a BLAST search of all 93 sequences would have yielded all alternative conformations. Furthermore, AF2 showed less success predicting fold switchers outside of its training set: 13% (1/8). This one success was a homolog of *E. coli* RfaH, a fold switcher with both conformations likely in AF2’s training set. Notably, AF2 failed to predict correct conformations of both new targets. It missed the 3-α-helical bundle conformation of an engineered protein that switches folds in response to temperature (28), and it predicted a conformation of the human protein isoform BCCIPα that differed completely from its experimentally determined structure (29). Since this structure is in complex with another protein, it is possible that BCCIPα may assume the AF2-predicted conformation in its uncharacterized apo state or in complex with a different binding partner. Nevertheless, neither AF2 nor AF-cluster predicted its experimentally characterized structure. These results suggest that current implementations of AF2 are unlikely to foster broad discovery of new fold switchers.

This study involved extensive sampling of fold-switching proteins, generating >450,000 structures for 100 fold switchers. Nevertheless, more sampling with more sophisticated techniques may enable AF2 to predict more alternative conformations not identified here. Though there is value in exploring and developing these techniques, our results indicate that AF2 is currently not sensitive enough to predict many new fold switchers from genomes.

AF2’s inability to accurately predict and score the multiple experimentally determined conformations of fold switchers suggests that the model has more to learn about protein energy landscapes (23). Complete understanding would enable AF2 to accurately predict both conformations of fold-switching proteins and their relative frequencies. AF2’s lack of understanding is evidenced by (1) its inability to predict >75% of fold switchers likely in its training set, (2) its inability to predict >87% of fold switchers outside of its training set, (3) its failure to accurately score models of alternative conformations, and (4) the inability of its structure module to distinguish between low and high energy conformations of fold-switching proteins. These findings are consistent with recent reports of unphysical and imbalanced AF2 predictions of experimentally determined protein kinase conformations (21). AF2 was trained mainly on single-fold proteins. Thus, its tendencies to predict one fold from a fold-switching sequence and inaccurate scoring of fold switchers likely reflect the limitations of what it has learned from its training set (9).

Some of AF2’s predictive unreliability appears to arise from faulty associations between sequence and structure. For instance, both AF2 and AF-cluster completely miss the experimentally determined conformation of BCCIPα, instead associating its sequence with the structure of BCCIPβ, a close homolog likely in AF2’s training set (29). Further, both AF2 and AF-cluster incorrectly predict only β-roll folds for CTDs of three fold-switching NusG proteins with ground state α-helical conformations. AF-cluster also incorrectly predicts that the β-roll CTDs of two single-folding NusG proteins can assume α-helical conformations indicative of fold switching. Thus, unlike its recently reported performance on some KaiB proteins (17), all of which were ≥ 47% identical to sequences of their PDB homologs, AF-cluster does not reliably associate sequence-diverse NusG proteins with their experimentally observed conformations.

Our results suggest a way to potentially improve AF2-based predictions of fold-switching proteins. Previous work from our lab shows that coevolutionary signals for both folds of fold-switching proteins are often present in MSAs (11). Deep MSAs show strong signal for a dominant conformation, while shallower subfamily-specific MSAs show increased signal for the alternative. AF2 misses this information in 44/56 cases. Better results may be obtained by fine-tuning AF2 to associate MSAs of different depths with different folds. AF-cluster’s success at predicting the structures of some KaiB homologs with shallow subfamily-specific MSAs indicates that such an approach has potential, and additional fine-tuning may strengthen the sequence-structure associations needed to predict alternative conformations of fold-switching proteins.

Nevertheless, deep learning models are limited by both their underlying assumptions and their training datasets. With very limited mechanistic understanding (23) and relatively few atomic resolution examples of fold switchers (10), it may not yet be possible to leverage deep learning to consistently predict this emerging phenomenon. There may be much about the protein universe– and particularly fold switching–that has not yet been observed. This “dark matter” is a new frontier of protein science.

## Methods

### The dataset

The dataset of fold-switching proteins having identical to high sequence similarity but assuming two distinct secondary/tertiary structures (folds) with experimentally determined structures (13) was used for the analysis (**Table S1**). Sequences of experimentally characterized RfaH/NusG variants (24) and two examples of folds-switching proteins identified in 2023 (28, 29) were also analyzed.

### Defining Fold1 and Fold2

Fold-switching proteins have two distinct conformations, A and B. Proteins with higher TM-scores in the fold-switching region for at least 3 out of 5 of their AlphaFold2.3.1 predictions were designated “Fold1” and the other conformation in the protein pair was denoted as “Fold2”, following the same ordering as in (13).

### AlphaFold2 (AF2) predictions

#### AF2.3.1 and AlphaFold-Multimer

The open-source version of AlphaFold2.2.0 and 2.3.1 (6) maintained on the NIH HPS Biowulf cluster (https://hpc.nih.gov/apps/alphafold2.html) was used to generate predictions. The template database contained PDB structures and sequences released till 2022-12-31. The pipeline was run both with and without templates, the predictions from the AlphaFold2-Multimer /AF2_multimer (7) pipeline were generated using both “monomer” and “multimer” option. **Table S1B** shows change in oligomeric state between the two folds/conformations.

#### AlphaFold2 with single sequences

Additional runs were performed using AF2.3.1 and AF2.2.0, with and without templates, simply putting in the target sequence in the prediction pipeline without generating MSA, to exclude any coevolutionary information that may be present in the MSA.

### Sampling prediction ensembles with AF2

#### Modified implementation of SPEACH_AF

Alanine-masked multiple sequence alignments (MSAs) were generated by identifying all amino acids in contact with a region of interest and mutating all contacting amino acids to alanine those within 4 residues of primary sequence to the region of interest (16). The region of interest was defined as a sliding window of 11 residues that moved by increments of 1 from the beginning to the end of the fold-switching region of each of 93 proteins. Positions in the MSA corresponding to residues within 4Å of any amino acid within a given region of interest–except those within 4 residues of primary sequence of that region–were converted to alanine except for the target sequence. Runs using the modified MSAs were carried out with AF2, with three random seeds for each MSA for a total of 15 models for each 11-residue window. A total of 77,160 predictions were generated using this method.

#### AF_Cluster

To perform more extensive sampling of conformations, AF-cluster was run with ColabFold (30) maintained on the NIH HPS Biowulf cluster (https://hpc.nih.gov/apps/colabfold.html). This module was used to generate multiple sequence alignments (MMseqs2-based routine (31)) for the proteins in our dataset using the UniProt database (32). The AF_Cluster pipeline (https://github.com/HWaymentSteele/AF_Cluster) (17) was then implemented to cluster the MSAs and these shallower MSAs were then used to generate predictions using ColabFold1.5.2 (which uses AF2.3.1) and ColabFold1.3 (utilizes AF2.2.0, to match the results presented in *Wayment-Steele et. al*, 11). The ColabFold1.3 run reproduced Wayment-Steele et al.’s predictions of both conformations of KaiB, RfaH, and Mad2. Both versions of ColabFold were run on all fold switchers, each generating 5 relaxed structures from two random seeds, 10 structures/shallow MSA, 3 recycles. Additionally, we ran ColabFold1.5.2 generating 50 relaxed models from 10 random seeds and 3 recycles on all NusG variants not in the PDB along with Sa1 and BCCIPα. Results for these variants outside of the PDB comprise all runs. Further, we repeated the 50-structure ColabFold1.5.2 runs with dropout and found no increase in alternative conformation sampling.

A table of total number of predictions generated for each protocol is presented in **Table S2**. All predictions following the AF_Cluster pipeline, were generated without templates, as in the original manuscript (17).

### Assessment of prediction quality

The per-residue Local Distance Difference Test (pLDDT) scores (a per-residue estimate of the prediction confidence on a scale from 0 – 100), quantified by determining the fraction of predicted Cα distances that lie within their expected intervals were used to determine confident predictions (33). The values correspond to the model’s predicted scores based on the lDDT-Cα metric, a local superposition-free score to assess the atomic displacements of the residues in the model. Values ≥ 90 were denoted as high confidence, and values between 70 to 90 are deemed confident.

Predictions were compared to the original experimentally determined structures using TM-align (18), (an algorithm for sequence-independent protein structure comparisons) and root mean square deviations (RMSDs) involving backbone atoms (C,Cα,N and O) calculated using biopython’s PDB.Superimposer module (34). TM-align first generates an optimized residue-to-residue alignment based on connections among secondary structural elements using dynamic programming iterations and then builds an optimal superposition of the two structures. TM-score (ranging from 0 to 1) is reported as the measure of overall accuracy of prediction for the models after the alignment, 0.6 signifying roughly similar folds for protein regions of interest. RMSD values ≤ 5 Å were used to infer similar structures.

### Reranking predictions based on pLDDT scores for AF_Cluster predictions

For an agnostic view of the pool of predictions generated for each protein, we reranked the predictions according to the percentage of confident residues (residues having pLDDT scores ≥70) and then compared them to the experimental structures. The predictions were designated as **Medium** (≥70% residues with pLDDT scores ≥ 70), **Good** (≥80% residues with pLDDT scores ≥70) and **High** (≥90% residues with pLDDT scores ≥70) confidence models. The predictions were rescored according to the percentage of confident residues in each pool of **Medium, Good, High** and **All** (includes all predictions for the protein) confidence models.

To determine which conformations were present among models within each of the four categories (**All, Medium, Good, and High**), the total number, N_ij_, of models corresponding to each conformation (i) of each fold-switching protein (j) were tabulated. A given conformation (ij) was considered to be predicted if N_ij_ ≥1.

### Prediction success

Success rate or prediction success is defined as the fraction of proteins for which at least one prediction corresponds well (TM-score of fold-switching region>0.6 (35)) to Fold1 or Fold2 (*Defining Fold1 and Fold2*). If the TM-scores for both Fold1 and Fold2 (TM-score1 and TM-score2, respectively) are greater than 0.6 the conformation is assigned to the conformation that produces the larger TM-score. The third label (other than Fold1 and Fold2) is “Other”, a.k.a. experimentally unobserved predictions, are designated to those predictions with TM-scores (TM-score1 and TM-score2) less than 0.6. After reranking, we checked for prediction success in Top1 (most confident prediction overall), Top10 (10 most confident predictions overall) and All (all predictions regardless of confidence) in the pool of predictions.

### AF2Rank

Starting with our original dataset (13), any proteins where one structure included only a short fragment or that had long gaps in the fold-switching region were excluded from the AF2Rank protocol. The final dataset consisted of 76 proteins (PDB IDs highlighted in supporting data, Table S1A).

Structures corresponding to each fold-switched conformation were passed to the AF2Rank protocol (7) as templates, and the candidate structure’s accuracy is assessed based on confidence scores of the AF2 output model. Before being passed to AlphaFold2, sidechain atoms were removed to prevent AF2 from using the underlying amino acid sequence to influence its prediction confidence. Beta carbons were added to glycine residues to mask their identity. AF2 was run without a MSA to remove coevolutionary influence from protein structure prediction. As in the original publication, a composite score of predicted local distance difference test (pLDDT), predicted template modeling score (pTM), and template modeling (TM) score was considered to be an energy function that evaluates model quality: the more confident and closer to the experimental structure, the higher the score (7). For each fold-switching protein, we passed AF2Rank each of the two folds as a template structure, using its amino acid sequence as the input sequence. pLDDT, pTM, and composite scores were compared between the two runs to determine which fold AF2 assigns higher confidence scores. TM-scores were also calculated between the output and template structures to assess prediction quality.

To ensure that we passed the same sequence to AF2 for fold-switched conformations, we truncated extraneous N- and C-terminal residues used for purification but endogenous to their respective sequences. If one structure included a domain that was not present in the other structure, that protein was excluded from the dataset. Any short gaps in the structures were modeled with RosettaCM (36), and the top scoring Rosetta model (minimum 1000 models generated) with a TM-score greater than 0.9 compared to the native structure were then selected for use. Hetero-atoms from non-standard residues such as the selenium in seleno-methionine and seleno-cysteine were replaced with their standard analogs (e.g. methionine and cysteine) using RosettaCM.

## Scripts and figures

The scripts used for all analyses were written in Python3; PyMOL (37) was used to visualize protein and plots were created using Matplotlib (38) and seaborn (39).

## Data sharing

Data and code used to generate the results in this manuscript can be found at: https://github.com/porterll/AF2_benchmark

## Acknowledgements

We thank Leslie Ronish, Samuel Lee, and Carolyn Ott for helpful discussions and Loren Looger, Marius Clore, and George Rose for commenting on the manuscript. This work utilized resources from the NIH HPS Biowulf cluster (http://hpc.nih.gov), and it was supported by the Intramural Research Program of the National Library of Medicine, National Institutes of Health (LM202011, L.L.P.).

