## Supplement for "AlphaFold2 has more to learn about protein energy landscapes"

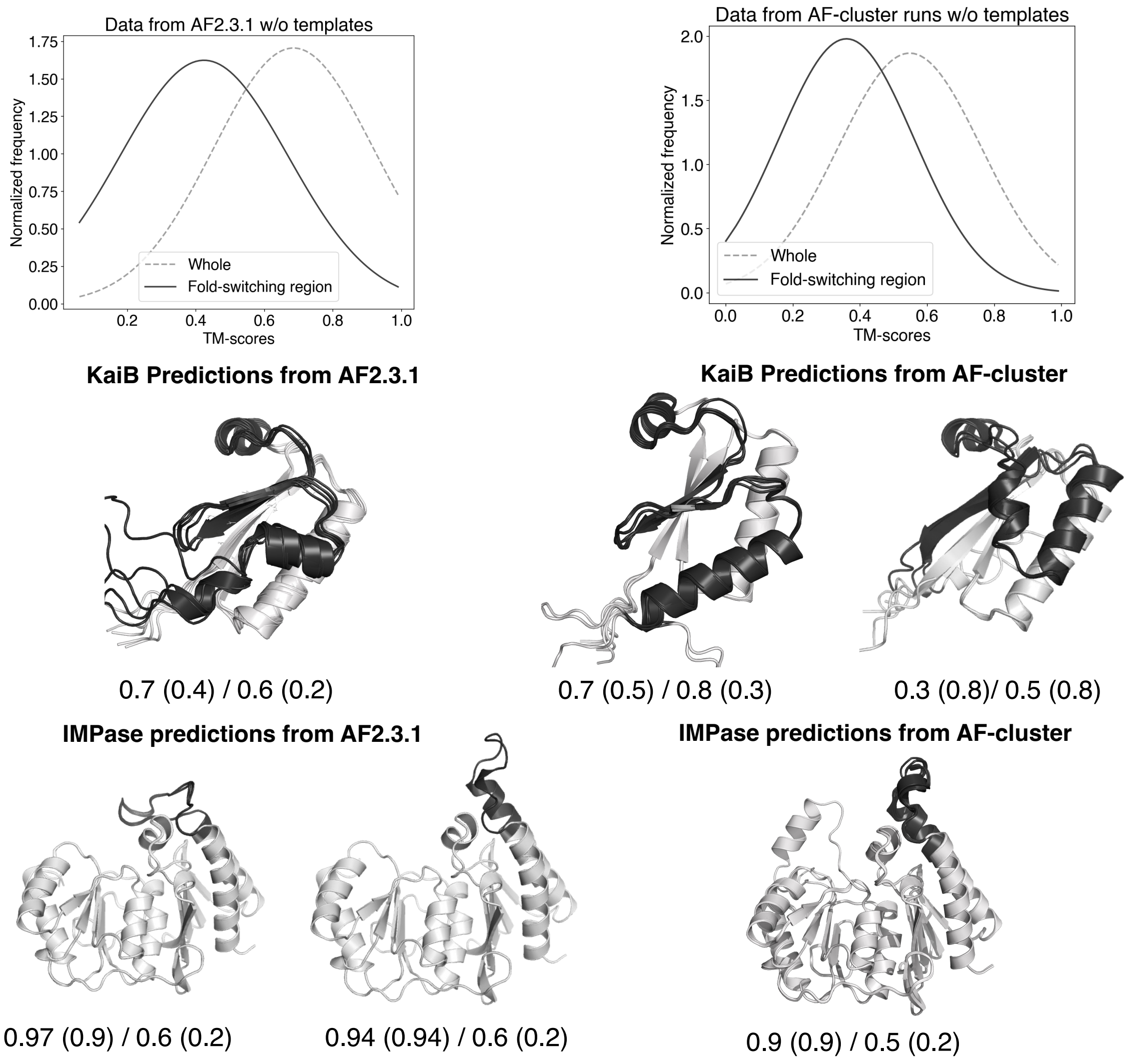


**Figure S1**. **TM-scores for fold-switching regions represent predictions of fold switchers more accurately than TM-scores of whole proteins**. Distributions of overall vs fold-switching region TM-scores for AF2.3.1 (upper left) and AF-cluster (upper right) demonstrate that whole-protein TM-scores overestimate prediction accuracies corresponding to regions of interest. Examples of predictions from AF2.3.1 and AF-clusters for KaiB and IMPase further demonstrate this point (fold-switching region is highlighted in black and the rest is grey). TM-scores relative to Fold 1 (left of /) and Fold 2 (right of /) are systematically higher for whole proteins (numbers without parentheses) compared to their fold-switching regions (in parentheses).

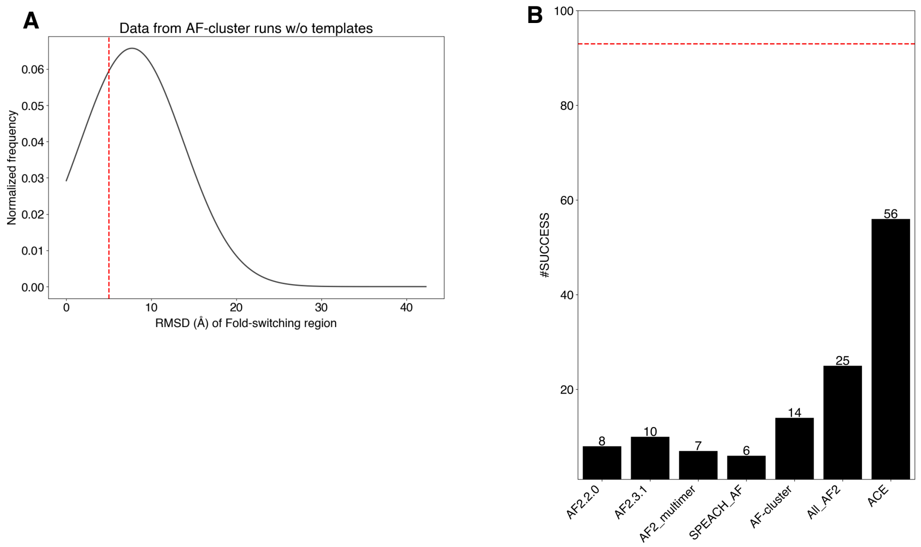


**Figure S2. Assessing predictions by RMSD yields results similar to TM-score based assessments**. Distribution of RMSD for the fold-switching region of AF-cluster predictions referenced against the fold-switching regions of the most similar experimentally determined structure is presented on the left panel (**A**), with threshold line at 5 Å. Prediction success measured by RMSD (**B**, fold-switching RMSD within 5 Å of experiment) yields results similar to TM-score (Figure 1 in main text).

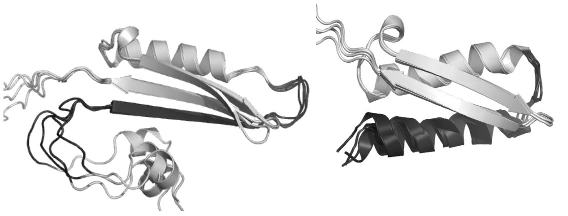

TM-scores (Whole/FS) - 0.6 (0.2) / 0.8 (0.4)

TM-scores (Whole/FS) - 0.5 (0.4) / 0.7 (0.4)

**Figure S3. AlphaFold2 is likely overtrained on some structures.** For example, Fold1 (PDBID: 2kxo) of MinE is predicted by inputting the full length MSA with templates into AF2.3.1, whereas Fold2 (PDBID:3r9j) is predicted by all models by inputting a single sequence (the sequence of MinE) with no templates. TM-scores relative to Whole/Fold-switching region (FS, black) are shown for full protein relative to experimentally determined structures (TM-scores relative to less similar fold in parentheses). Regions of the protein that don't switch folds are light gray.

A

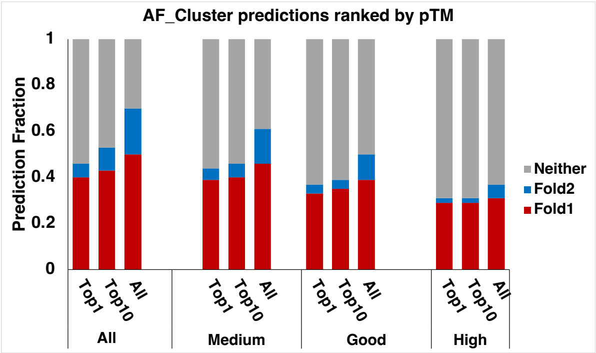


B

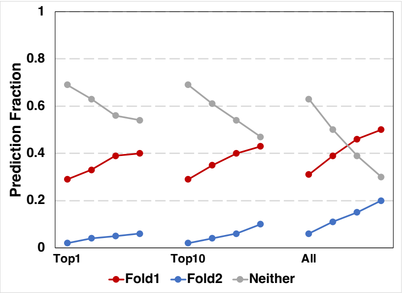


**Figure S4. Ranking by predicted template modeling (pTM) score selects against experimentally observed conformations in favor of experimentally unobserved for AF-cluster predictions.** (A) Bar-plot representation of the Prediction Fraction in Top1,Top10 and All ranked models. (B) Trendline plots showing the change in prediction success in categories –High (pTM>0.9), Good (pTM>0.7), Medium (pTM > 0.6), and All, respectively for Top1, Top10 and All predictions. Neither denotes predictions whose fold-switching regions had TM-scores < 0.6 relative to the experimentally determined structures of both Fold1 and Fold2.

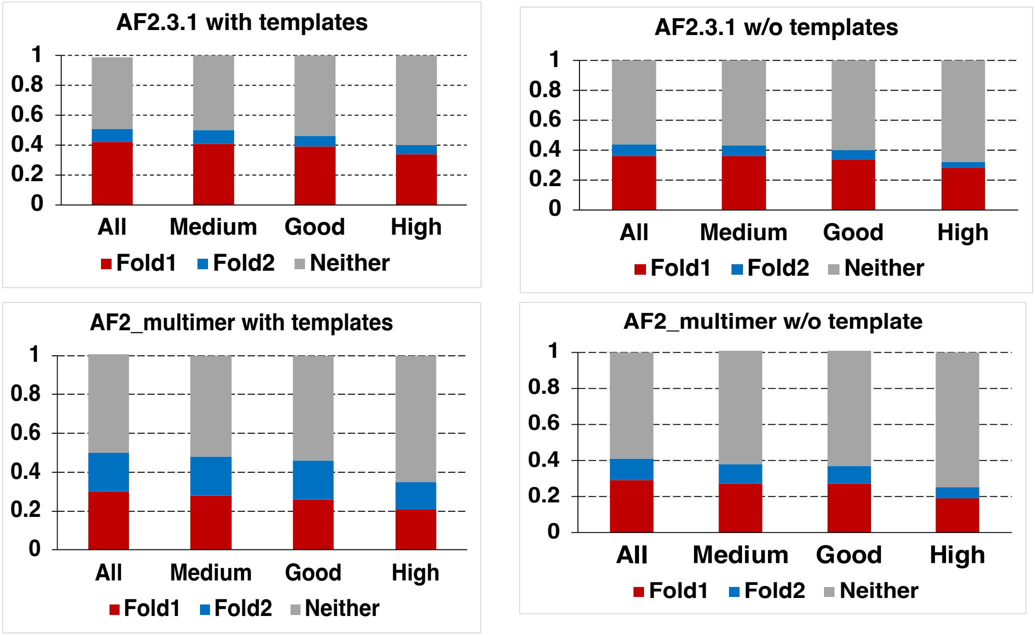


**Figure S5. pLDDT scores select against experimentally determined conformations of fold-switching regions in all AF2.3.1. runs.** Predictions are ranked by confidence (percentage of residues with pLDDT scores > 70). The categories are defined as - All, Medium (confidence > 70%), Good (confidence > 80%) and High (confidence >90%). Neither denotes predictions whose fold-switching regions had TM-scores < 0.6 relative to the experimentally determined structures of both Fold1 and Fold2.

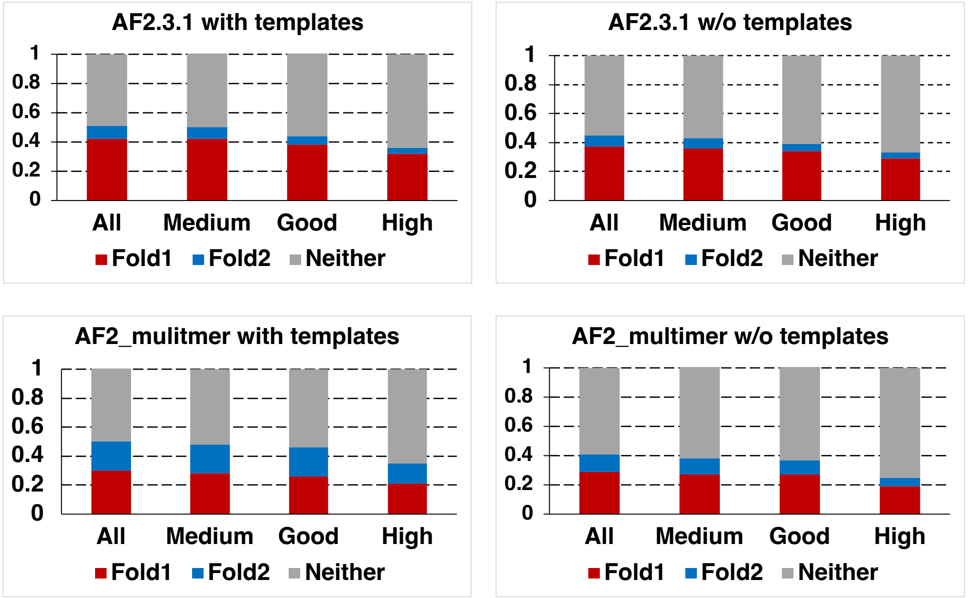


**Figure S6. pTM scores select against experimentally determined conformations of fold-switching regions in all AF2.3.1. runs.** Predictions are ranked by confidence (pTM score defined as - All, Medium (pTM ≥ 0.6), Good (pTM ≥ 0.7) and High (pTM ≥ 0.8). Neither denotes predictions whose fold-switching regions had TM-scores < 0.6 relative to the experimentally determined structures of both Fold1 and Fold2.

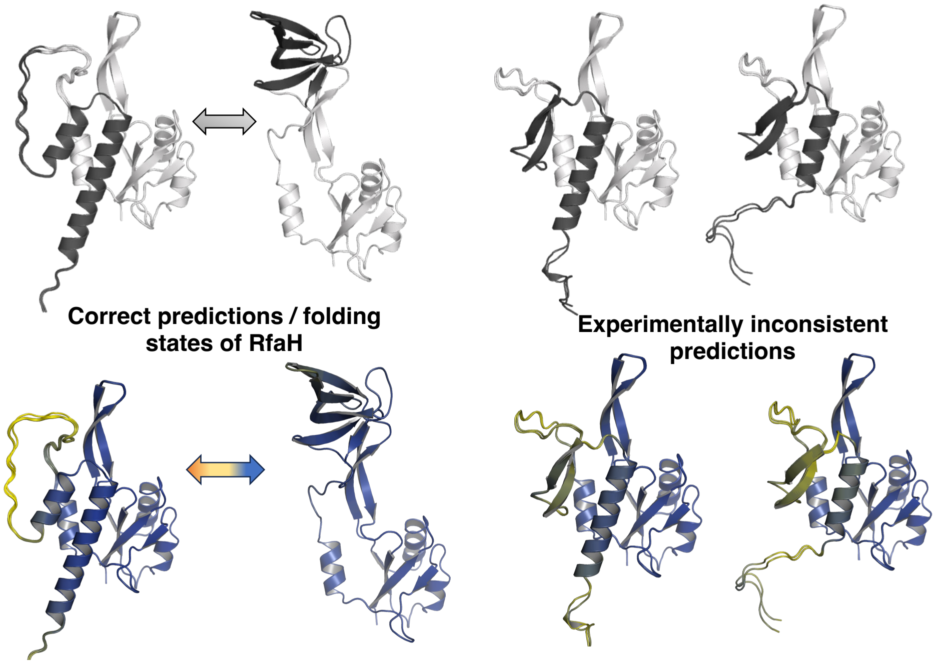


**Figure S7. AF2 predicts experimentally inconsistent conformations of RfaH.** Models corresponding to experimentally determined structures on left; experimentally inconsistent predictions shown on right. The figures below are colored by pLDDT scores, (color ranging from orange, yellow to blue, corresponding to pLDDT scores from 0 to 100). All predictions were generated from AF2_multimer (run without partner and no templates).

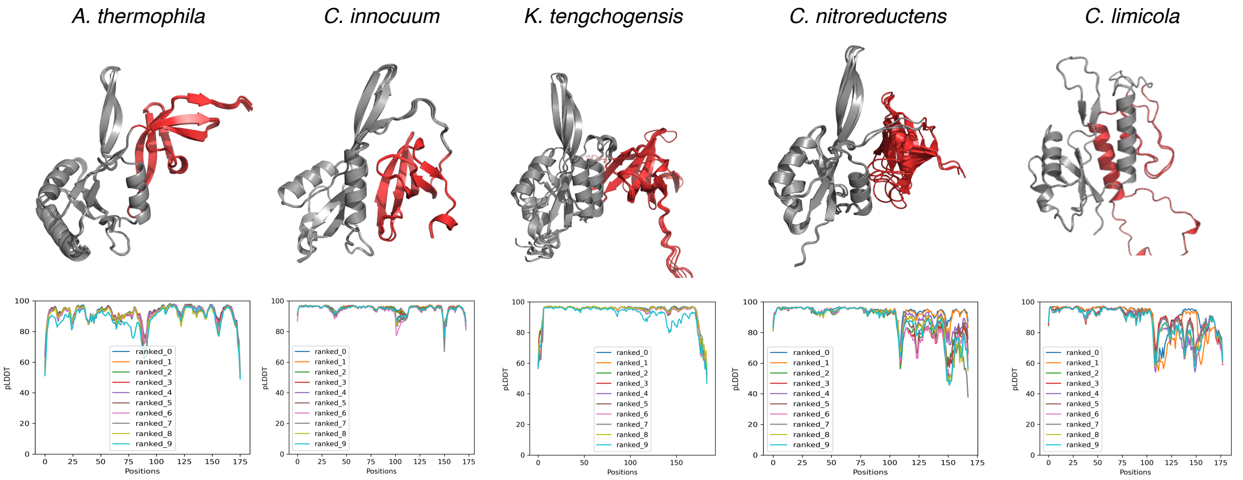


**Figure S8. AlphaFold2.3.1 fails to predict the experimentally confirmed helical conformations in the C-terminal domains (CTDs) of 4/5 RfaH homologs.** In all variants but *C. limicola*, only β-sheet CTDs (red) are predicted. Structurally conserved N-terminal domains are colored gray. pLDDT scores of all 10 models of each protein generated without templates are shown below their predicted structures. C-terminal domains comprise residues 115-end of protein.

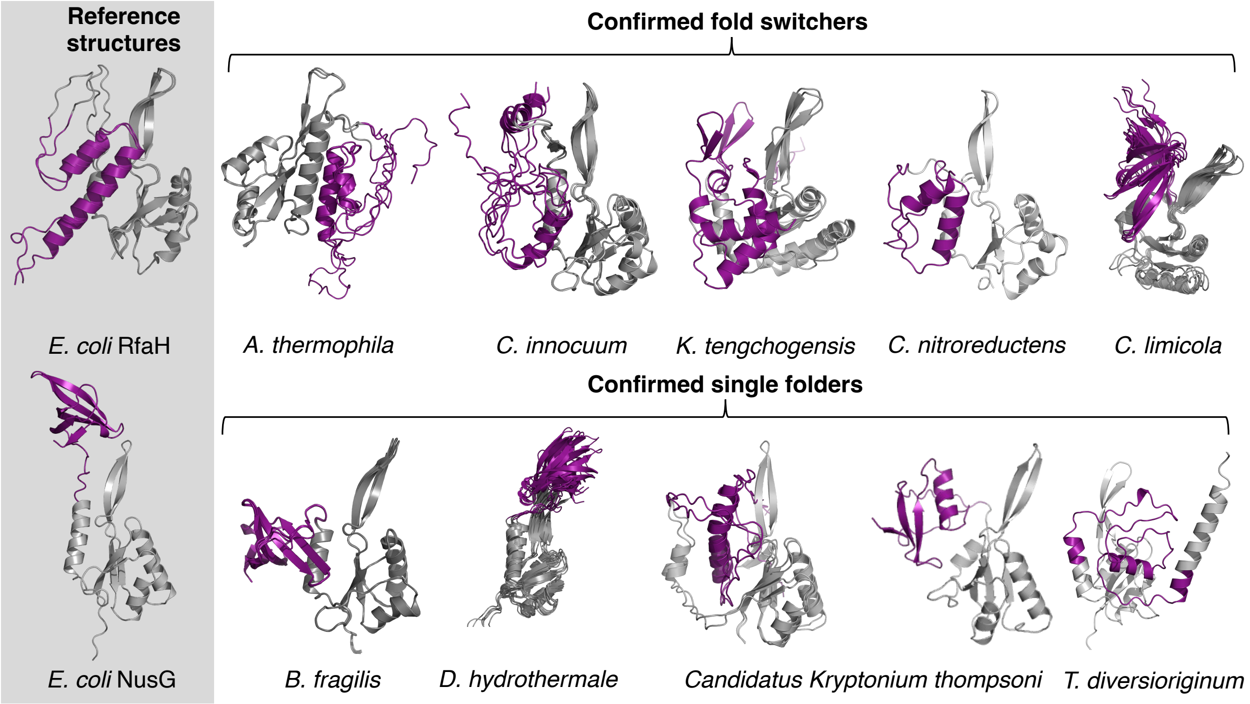


**Figure S9. AF-cluster predictions cannot distinguish between RfaH homologs with helical C-terminal domains (CTDs, upper row) and β-sheet C-terminal domains (lower row).** Further, pLDDT scores of all helical CTD predictions are low (average ≤50), further indicating that correct and incorrect predictions cannot be distinguished. All CTDs are colored purple; structurally conserved N-terminal domains are gray. Experimentally confirmed reference structures of *E. coli* RfaH and NusG are shown on the left column with the same color scheme.

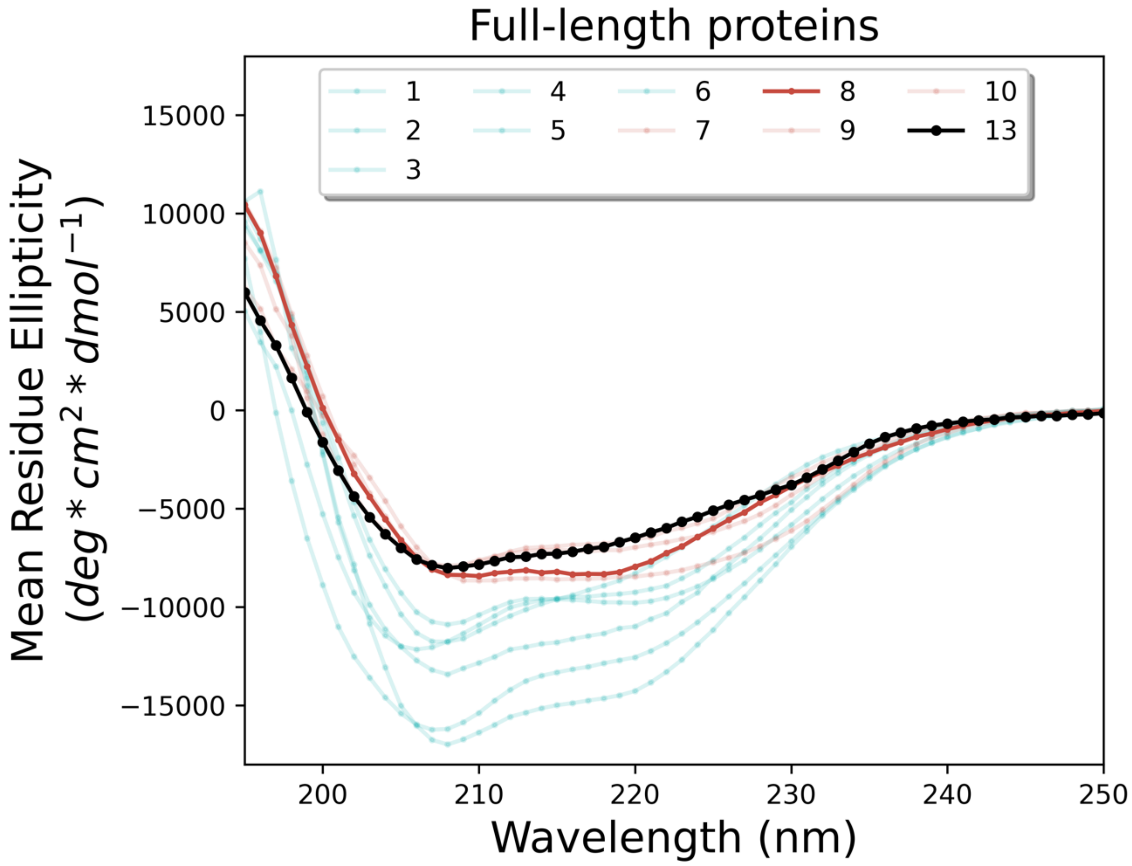


**Figure S10. The circular dichroism spectrum of NusG Variant 13 (black) resembles single-folding NusGs with ground state β-sheet folds (red) rather than fold-switching NusGs with ground state α-helical folds (teal)**. CD spectra of all variants except for 13 were taken from reference 24 in the main text. AF-cluster predicted that it and Variant 8 (bold red) can assume helical folds, inconsistent with experimental evidence. The sequence of Variant 13 (Table S4) inserted into a pET-28a(+) vector was purchased through BioBasic, codon optimized for *E. coli*. Variant 13 was purified using Cytiva Hi-TRAP columns on an ÄKTA Pure at room temperature. Its 6x-His tag was cleaved overnight with biotinylated thrombin (Sigma Millipore) at 4°C while dialyzing in 100 mM potassium phosphate, 10% glycerol (v/v) pH 7.4 using a ThermoFisher dialysis cassette (10 kDa MWCO). The cleaved sample was again run on a Hi-TRAP column, and the unbound flow-through was then concentrated in a Millipore centrifugal concentrator (10 kDa MWCO) and subsequently polished through size exclusion chromatography with a Superdex 70 Increase 10/300 column (Cytiva) and was found to be monomeric. Its CD spectrum was collected on a Chirascan spectrometer (Advanced Photophysics) in 100 mM Phosphate, pH 7.6 at 9µM.

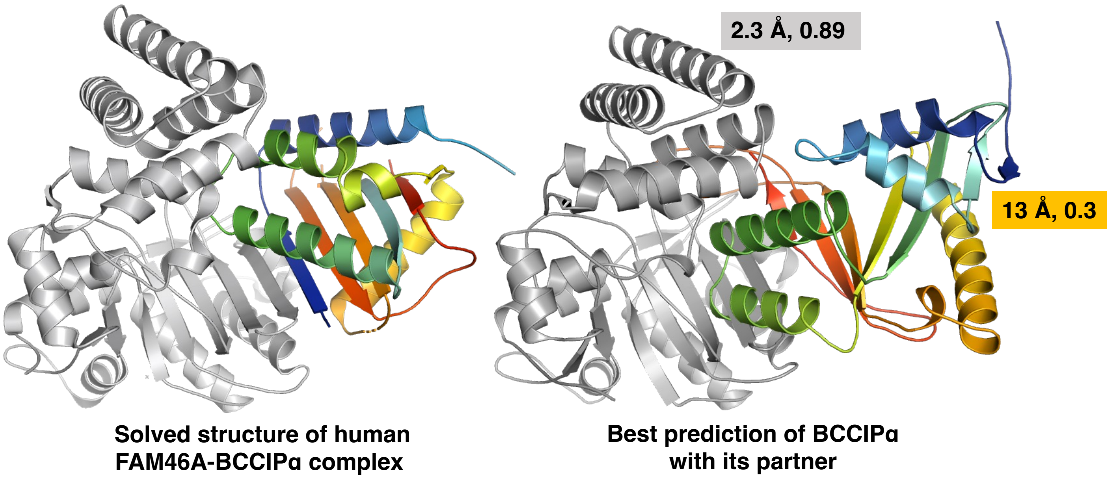


**Figure S11. Predicting the structure of BCCIPα with its binding partner using AF2_multimer generated the incorrect BCCIPβ conformation for all models.** Structure of the experimentally determined complex (PDB ID: 8EXF, left) differs from all AF2_multimer models (Ranked 1 shown). The structure of BCCIPα's binding partner, FAM46A (gray), was predicted with has high accuracy: TM-score 0.89 and RMSD 2.3 Å. Whereas BCCIPα (rainbow N->C, blue->red) was poorly predicted: TM-score 0.3 and RMSD 13 Å. The predicted binding interface is also incorrect.
